# Antigenic Divergence of Cobra Short-chain α-Neurotoxins: Implications for Regional Antivenom Effectiveness in Southeast Asia

**DOI:** 10.64898/2025.12.15.694306

**Authors:** Choo Hock Tan, Praneetha Palasuberniam, Lee Louisa Pernee, Kae Yi Tan

**Affiliations:** School of Medicine, College of Life Sciences and Medicine, National Tsing Hua University, Hsinchu, Taiwan; Institute of Bioinformatics and Structural Biology, College of Life Sciences and Medicine, National Tsing Hua University, Hsinchu, Taiwan; Venom Research & Toxicology Research Lab, Department of Pharmacology, Faculty of Medicine, University of Malaya, Kuala Lumpur, Malaysia; Department of Biomedical Sciences, Faculty of Medicine and Health Sciences, Universiti Malaysia Sabah, Kota Kinabalu, Sabah, Malaysia; Protein and Interactomics Lab, Department of Molecular Medicine, Faculty of Medicine, University of Malaya, Kuala Lumpur, Malaysia

**Keywords:** Antivenom, Immunoreactivity, Snakebite envenoming, Cobra venom

## Abstract

Short-chain α-neurotoxins (SNTXs) are the principal neurotoxic components in several Asiatic cobras, including *Naja atra*, *Naja philippinensis*, and *N. samarensis*. Although structurally conserved, SNTXs exhibit marked antigenic variation that can limit the effectiveness of regional antivenoms used for snakebite envenoming in Asia. Here, we evaluated the immunoreactivity of the Philippine Cobra Antivenom (PCAV) and three regional products—*Naja kaouthia* Monovalent Antivenom (NkMAV), Neuro Bivalent Antivenom (NBAV), and Indonesian SABU—against a panel of cobra venoms and purified α-neurotoxins. PCAV bound strongly to homologous *N. philippinensis* SNTX but showed weak cross-reactivity with SNTXs from *N. kaouthia*, *N. sputatrix*, and *N. atra*, and minimal recognition of marine elapid SNTXs and long-chain α-neurotoxins (LNTXs) of Monocled Cobra as well as King Cobra. Conversely, NkMAV and NBAV recognized mainland Asian SNTXs more broadly but reacted poorly with the Philippine cobra toxin. Hierarchical clustering of normalized immunoreactivity delineated two major SNTX antigenic subtypes corresponding to Philippine versus continental Asian lineages. Peptide sequence analysis unmasked two distinct loop-II motifs (^28^WWS–TII^37^ and ^28^RWR–YRT^37^) associated with these divergent immunotypes. Phylogenetic reconstruction suggests that the Philippine cobras retain an ancestral SNTX motif, while Sundaic and East Asian cobras have diversified to employ another major SNTX form. Epitope prediction further revealed differences in surface exposure and accessibility that help explain the limited cross-neutralization across antivenoms. These findings demonstrate a clear antigenic dichotomy among Asian cobra SNTXs, which underlies species-specific antivenom effectiveness and highlights the need to incorporate representative SNTX variants into immunogen formulations to improve regional antivenom coverage.

## Introduction

Snakebite envenoming is a WHO-listed priority neglected tropical disease that disproportionately affects rural and low-income communities across Asia [1]. As a One Health challenge spanning human health, biodiversity, and regional healthcare inequities, improving antivenom coverage requires a deeper understanding of venom variation among medically important snakes [2, 3]. Cobra venoms, in particular, exhibit striking interspecific and geographical diversity in toxin composition, which directly influences clinical severity and treatment outcomes [3]. Within these venoms, three-fingered protein toxins represent the dominant neurotoxic components responsible for rapid paralysis in neurotoxic snakebites. Proteins with the three-finger fold are widely distributed among metazoans and play critical physiological roles. For instance, the GPI-anchored Lynx1 protein modulates cholinergic transmission by fine-tuning nicotinic acetylcholine receptors (nAChRs) in mammals, influencing learning, memory, and neural plasticity. Similarly, secretory three-finger proteins (TFPs) such as SLURP-1 and SLURP-2 are involved in cell proliferation, immune responses, and tissue repair [4–6]. In some advanced snakes, these ancestral physiological proteins were co-opted for neofunctionalization—the inherent plasticity of the three-finger motif has allowed it to be exploited and repurposed for diverse toxic effects, ranging from cytotoxicity to neurotoxicity [7, 8]. This evolutionary adaptation has given rise to a new class of TFPs known as three-finger toxins (3FTXs), which are predominantly found in the Elapidae family, including cobras, kraits, king cobras, coral snakes, mambas, and sea snakes [3, 9–11].

Within the 3FTX family, alpha-neurotoxins are among the most well-studied members. These toxins act as high-affinity competitive blockers of postsynaptic muscle-type nAChRs [12]. The alpha-neurotoxins are broadly classified into two subfamilies based on peptide length and the number of intramolecular disulfide bonds: (1) short-chain α-neurotoxins (SNTX), which have 60–62 residues and four disulfide bridges; and (2) long-chain α-neurotoxins (LNTX), which consist of 66–75 residues and five disulfide bridges. In cobras (*Naja* spp.), alpha-neurotoxins are the principal 3FTXs responsible for neurotoxic envenomation, resulting in systemic paralysis and respiratory failure as the mode of lethality. The abundance of these toxins in venom correlates positively with venom lethality, as measured by intravenous median lethal doses (LD_50_), where higher levels of alpha-neurotoxin protein correspond to greater venom potency [12].

The distribution of alpha-neurotoxins in cobra venom varies across species and geographical regions. Recent findings from cobra venomics and phylogeographical studies reveal a phenotypic dichotomy in venom composition, characterized by a transition in alpha-neurotoxin subtype that accompanies the biogeographical distribution of cobra species, honed by evolutionary pressures to fulfil specific ecological roles. In *Naja naja* (Indian cobra), LNTX predominates, while in Asiatic cobras that have dispersed eastward, a more evolutionarily derived form—without loss of lethality—has largely replaced LNTX. For example, venoms of *N. atra* (Taiwan), *N. kaouthia* (Vietnam), *N. philippinensis*, and *N. samarensis* (Philippines) contain predominantly SNTX, with LNTX virtually absent [12–15]. Despite their structural similarities, SNTX and LNTX exhibit distinct pharmacological profiles at nicotinic acetylcholine receptor (nAChR) subtypes. LNTX is often considered more medically relevant in snakebite envenomation due to its stronger binding affinity toward human nAChRs compared to SNTX [16, 17]. This increased affinity is likely due to the additional disulfide bond in loop II of LNTX, which SNTX lacks. Whether this translates to reduced fatality in human envenomation is inconclusive, and the toxicity of SNTX should be reckoned with for its high potency and rapid action. Of note, both SNTX and LNTX have comparable lethality in rodent models (LD_50_ values typically range between 0.1 and 0.2 µg/g, intravenously), and the lethality of venom during envenomation depends on the abundance of the dominant alpha-neurotoxin form. For instance, Philippine cobras (*N. philippinensis*) cause prominent neurotoxicity with rapid onset in humans, despite SNTX being the only alpha-neurotoxin present in their venom (Watt et al., 1988).

From the treatment perspective, clinically effective antivenoms must be able to immunorecognize and neutralize the dominant neurotoxin form. However, antivenoms often exhibit lower neutralization activity against SNTX than LNTX, possibly due to the limited antigenicity of SNTX proteins [18–20]. SNTX and LNTX share common epitopes within their respective subgroups, enabling cross-reactivity and cross-neutralization by antivenoms raised against different cobra species in Southeast Asia. For instance, *N. kaouthia* Monovalent Antivenom (NkMAV) produced in Thailand can cross-neutralize the venoms of *N. sumatrana* (Equatorial Spitting Cobra, Malaysia) and *N. sputatrix* (Javan Spitting Cobra) [19, 21]. However, this is not the case for *N. philippinensis*, as its venom toxicity can only be effectively neutralized by species-specific Philippine Cobra Antivenom (PCAV), not NkMAV [22]. Furthermore, a preliminary test with antivenom raised against *N. atra* (Neuro Bivalent Antivenom, Taiwan) has also shown ineffectiveness in neutralizing *the lethality of N. philippinensis venom* (unpublished). The finding was rather unexpected, as proteomic analyses have shown that *N. philippinensis* and *N. atra*, both considered the most eastern-dispersed cobras, possess only SNTX as their alpha-neurotoxins, albeit in differing abundances [12, 14, 23]. These observations raise a broader question of public health importance: Do antigenically distinct “immunoforms” of SNTX exist across Asian cobras, and could this divergence undermine the regional effectiveness of antivenoms used to treat neurotoxic envenoming? Understanding antigenic divergence is directly relevant to snakebite envenoming, a WHO-listed priority neglected tropical disease disproportionately affecting rural communities in Southeast Asia. Regional antivenoms are often relied upon across species and geographical boundaries; however, antigenic incompatibilities may limit cross-neutralization and contribute to treatment failures. While venom composition and sequence variation in cobra toxins have been studied extensively, the degree to which homologous α-neurotoxins differ immunologically remains unclear, particularly for SNTXs, which dominate in Southeast and East Asian cobras.

This study therefore evaluates the immunoreactivity of multiple regional antivenoms toward a panel of elapid venoms and purified α-neurotoxins. We focused on identifying patterns of SNTX antigenicity across species, determining whether distinct immunotypes exist, and characterizing sequence motifs that may underpin divergent antibody recognition. By integrating immunoprofiling with phylogenetic and sequence analyses, we aim to clarify how antigenic divergence among cobra SNTXs shapes antivenom cross-reactivity. These insights may guide the future formulation of antivenoms with broader regional utility and improved therapeutic coverage for neurotoxic snakebite envenomation.

## 2. Materials and methods

### 2.1 Venoms and antivenoms

Venoms of *Naja philippinensis* (Philippines)*, Naja sputatrix* (Indonesia), *Naja naja* (Pakistan), and *Naja kaouthia* (Thailand) were supplied by Latoxan (Valence, France). Venoms of *Naja sumatrana*, *Ophiophagus bungarus* (formerly *Ophiophagus hannah*), *Hydrophis curtus*, and *Hydrophis schistosus* were collected by author (CHT). *Naja atra* venom (Taiwan) was provided by Professor Inn-Ho Tsai from Academia Sinica, Taiwan while *Laticauda colubrina* venom was from Venom Supplies, Australia. *Calloselasma rhodostoma* venom was provided by the Queen Saovabha Memorial Institute (Bangkok, Thailand). All venom samples were stored in lyophilized form at −20 °C until use.

The isolation of alpha-neurotoxins from different venoms was based on chromatographic methods, as reported or established in previous proteomic studies by the same group. These include SNTXs from *N. philippinensis* [12], *N. sputatrix* [19], *N. sumatrana* [24], *N. kaouthia* [15], *Hydrophis curtus* [25], *Hydrophis schistosus* [10], *Laticauda colubrina* [26], and *N. atra* (unpublished). LNTXs were also isolated from venoms of *N. kaouthia* [15], and *Ophiophagus bungarus* (unpublished), respectively, for immunoreactivity study.

Philippine Cobra Antivenom (PCAV; Lot# 201804) used in the immunoreactivity is monovalent antivenom raised against *N. philippinensis* and is produced by the Research Institute for Tropical Medicine in the Philippines. *Naja kaouthia* Monovalent Antivenom (NkMAV; batch no.: NK00116) is a monospecific antivenom raised against Thai *Naja kaouthia* venom, produced by the Queen Saovabha Memorial Institute (QSMI), Bangkok, Thailand. Neuro Bivalent Antivenom (NBAV; batch no.: 61-06-0002) is a bivalent antivenom raised against Taiwanese *Naja atra* and *Bungarus multicintus,* manufactured by Central for Disease Control, Taipei, Taiwan. Serum Anti Bisa Ular (SABU; batch no.: 4701516) is a polyvalent antivenom raised against *Naja sputatrix* (Javan Spitting Cobra), *Bungarus fasciatus* (Banded Krait) and *Calloselasma rhodostoma* (Malayan Pit Viper), manufactured by BioFarma Pharmaceuticals, Bandung, Indonesia.

### 2.1 Determination of antivenom, venom and isolated toxin concentrations

The protein concentrations of venoms and isolated toxins were estimated by Nanodrop Spectrophotometer 2000/2000c (ThermoFisher™, Rockford, IL, USA). Venoms and isolated toxins were prepared at a stock concentration of 1 mg/ml for subsequent assays. Protein concentration of Philippine Cobra Antivenom (PCAV), *Naja kaouthia* Monovalent Antivenom (NkMAV), Neuro Bivalent Antivenom (NBAV) and Serum Anti Bisa Ular (SABU) were determined using the Thermo Scientific^TM^ BCA (bicinchoninic acid) Protein Assay Kit (Thermo Scientific^TM^ Pierce^TM^, Waltham, MA, USA). Bovine serum albumin (BSA) was used to obtain a standard curve, according to the manufacturer’s protocol. Antivenom concentrations of samples were determined in three independent experiments, and absorbance values were expressed as means ± S.E.M. The concentrations of PCAV, NkMAV, NBAV and SABU were determined to be 16.2 mg/ml, 57.54 mg/ml, 22.91 mg/ml and 238.76 mg/ml, respectively. Antivenom concentrations were further prepared at a stock concentration of 10 mg/ml and subsequently diluted to 1:450, standardized based on Tan *et al*. [22].

### 2.2 Antivenom immunoreactivity study

The immunoreactivity of Philippine Cobra Antivenom (PCAV), *Naja kaouthia* Monovalent Antivenom (NkMAV), Neuro Bivalent Antivenom (NBAV) and Serum Anti Bisa Ular (SABU) were examined with indirect enzyme-linked immunosorbent assay (ELISA) for the above-stated venoms and isolated toxins. For PCAV immunoreactivity, venoms of *Naja philippinensis* and *Calloselasma rhodostoma* were used as the positive and negative controls, respectively. Positive controls for NkMAV, NBAV and SABU were respective homologous venoms, *Naja kaouthia*, *Naja atra* and *Naja sputatrix*. *Trimeresurus albolabris* venom was uniformly used as a negative control for these three antivenoms.

In the ELISA experiment, the immunoplate wells were first coated with 10 ng of the respective venoms and isolated toxins overnight at 4 °C. The excess solution was removed by rinsing the wells four times with washing buffer. 100 μl of antivenoms (PCAV, NkMAV, NBAV or SABU) at a dilution of 1:450 were added into the venom- or NTX-coated wells, followed by incubation for 1 hr at room temperature. The wells were washed four times with washing buffer, flicked dry, followed by addition of horseradish peroxidase (HRP)-tagged secondary antibody (anti-horse HRP-IgG, 1:10000 (Jackson ImmunoResearch Inc. (West Grove, PA, USA) to each well. The mixtures were then incubated at room temperature for another hour and washed four times with wash buffer before adding 50 μl of TMB (3, 3’, 5, 5’-tetramethylbenzidine) (Merck, Kenilworth, NJ, USA), as the substrate. The enzymatic reaction was allowed to proceed for 25 min at room temperature (25 °C) in the dark, and subsequently terminated with 50 μL of 0.2 M sulfuric acid. The signals developed were read as absorbance at 450 nm using a Tecan M1000Pro Multimode plate reader (TECAN, Switzerland). The immunological binding activities of antivenom to venoms and isolated alpha-toxins were interpreted by absorbance unit at 450 nm. The assays were performed in three independent experiments, and absorbance values were expressed as means ± S.E.M.

### 2.3 Heatmap analysis

Mean ELISA absorbance values (A₄₅₀) obtained for each antivenom–toxin and antivenom–venom combination were normalized by row z-score transformation and visualized as clustered heatmaps using Perseus (version 2.1.5.0). Clustering of both rows (antivenoms) and columns (venoms or isolated α-neurotoxins) was performed using Euclidean distance and average linkage. The red–white–blue color scale represents relative reactivity, with red indicating stronger and blue weaker binding compared with the mean response of each antivenom.

### 2.4 Statistical analysis

Values were expressed as means ± SEM of three independent experiments. Differences between absorbance values of isolated alpha-neurotoxins were statistically evaluated using one-way ANOVA with Bonferroni’s *post hoc* test. Results were statistically significant at *p*-value < 0.05.

### 2.5 Multiple sequence alignment

Jalview software (version 2.11.1.3)[27] and the MUSCLE (Multiple Sequence Comparison by Log-Expectation) [28] tool were used to align multiple sequences. Multiple sequence alignment was performed using sequences from related species obtained from the UniProtKB depository (http://www.uniprot.org/) and from transcriptomics reported earlier [29, 30]. The sequences were chosen based on their medical relevance and phylogenetic relatedness of the species.

### 2.6 Phylogeny analysis

Sequences of short-chain alpha-neurotoxins annotated for Asian elapids (retrieved from Universal Protein Knowledgebase (UniProtKB) (accessed on: January 18, 2025) and transcriptome of *N. sumatrana* [29] were retrieved to construct a phylogenetic tree. The tree was reconstructed using [31]_maximum likelihood analysis with bootstrapping (100 replications) in the advanced mode of the phylogeny.fr web server (http://www.phylogene.fr/ (accessed on 20 January 2025), incorporating MUSCLE (v3.7) for multiple alignments, Gblocks (v0.91b) for alignment curation, and PhyML (v3.0) for tree building. Graphical representation and edition of the tree were performed using TreeDyn (v198.3).

## 3. Results

### 3.1 Immunoreactivity of PCAV toward Southeast Asian elapid venoms and isolated alpha-neurotoxins

PCAV showed varying immunoreactivity toward different short-chain and/or long-chain alpha-neurotoxins (SNTX and LNTX, respectively) isolated from the venoms of four Asian cobras (*N. philippinensis, N. kaouthia, N. sputatrix, and N. atra*), sea snakes (*H. curtus and H. schistocus*), sea krait (*L. colubrina*) and the King Cobra (*O. bungarus*) (Fig 1). The highest immunoreactivity was observed in *N. philippinensis* venom (positive control) and its SNTX (abs: 2.7) while its immunoreactivity toward *C. rhodostoma* venom (negative control) was negligible. The immunoreactivity of PCAV was noticeably reduced toward the SNTXs of other Asiatic cobras in the following order: *N. kaouthia* (abs: 2.2) > *N. sputatrix* (abs: 1.7) > *N. atra* (abs: 1.4). The immunoreactivity was low toward *L. colubrina* SNTX (abs: 0.5), and negligible toward the SNTXs of *H. curtus* and *H. schistocus* (abs: <0.1). PCAV also lacked immunoreactivity toward the LNTXs of *N. kaouthia* and *O. hannah* (abs: < 0.1). Statistically, the mean values of isolated SNTX and LNTX immunoreactivity from all species were significantly lower compared to SNTX from *Naja philippinensis* (one-way ANOVA, Bonferroni’s *post hoc* test, *p* < 0.0001)

**Figure 1.**
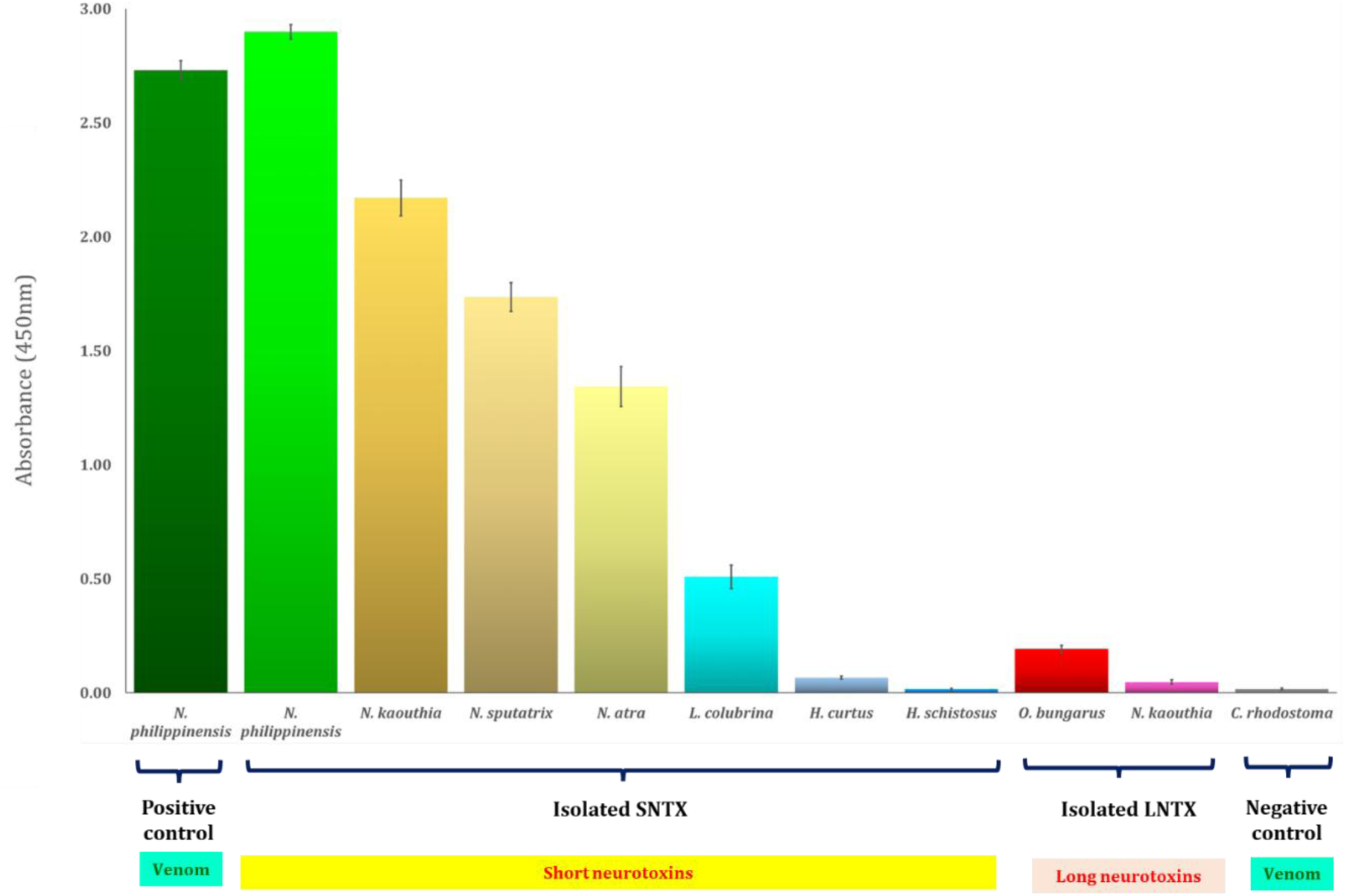
Immunoreactivity of Philippine Cobra Antivenom (PCAV) toward isolated alpha-neurotoxins (short-chain and long-chain alpha-neurotoxins) of Asian elapids. Abbreviation of genera: *N*: *Naja*, for true cobra; *H*: *Hydrophis*, for sea snake; *L*: *Laticauda*, for sea krait; *O*: *Ophiophagus*, for King Cobra; *C*: *Calloselasma*, for Malayan Pit Viper. Values were expressed as means ± SEM from three independent experiments. *N. philippinensis* and *C. rhodostoma* venoms were positive and negative controls for the assay.

### 3.2 Immunoreactivity of regional antivenom products toward Southeast Asian elapid venoms and isolated alpha-neurotoxins

Figure 2 shows the comparative immunoreactivity of NkMAV, NBAV and SABU toward the three homologous cobra venoms (*N. kaouthia*, *N. atra*, and *N. sputatrix*, respectively), and alpha-neurotoxins isolated from different elapid species as profiled in Figure 1. NkMAV, raised against the Thai Monocled Cobra, and NBAV, raised against the Taiwan Cobra and Taiwan Krait, showed relatively high immunoreactivity toward the venoms of *N. kaouthia*, *N. atra* and *N. sputatrix* (Abs: 3.0). SABU, raised against the Javan Spitting Cobra, Banded Krait and Malayan Pit Viper, had significantly lower immunoreactivity (Abs: 1.8-2.0) toward the three cobra venoms (including its own homologous *N. sputatrix* venom) compared to NkMAV and NBAV. Tested against isolated SNTX, NkMAV showed significantly lower immunoreactivity (Abs: 1.8) than NBAV (Abs: 2.5-3.0) toward the SNTXs of *N. kaouthia* and *N. sputatrix*, while both antivenoms show comparable immunoreactivity toward the SNTX of *N. atra* (Abs: 1.8). SABU showed a statistically much lower immunoreactivity than NkMAV and NBAV toward these SNTX of *N. kaouthia*, *N. atra*, and *N. sputatrix* (Abs: 0.4-0.8) (*p* < 0.001). Of note, NkMAV, NBAV and SABU all exhibited extremely weak immunological binding to SNTX of *N. philippinensis* and marine elapids (*H. curtus*, *H. schistosus*, *L. colubrina*) (Abs < 0.5), which is noticeably insignificant in comparison to the absorbance for the negative control, *T. albolabris* venom (Abs < 0.5). NkMAV was significantly more immunoreactive than NBAV and SABU toward its homologous LNTX (isolated from *N. kaouthia* venom), whereas all three antivenoms were weak in binding to the LNTX of the King Cobra.

**Figure 2.**
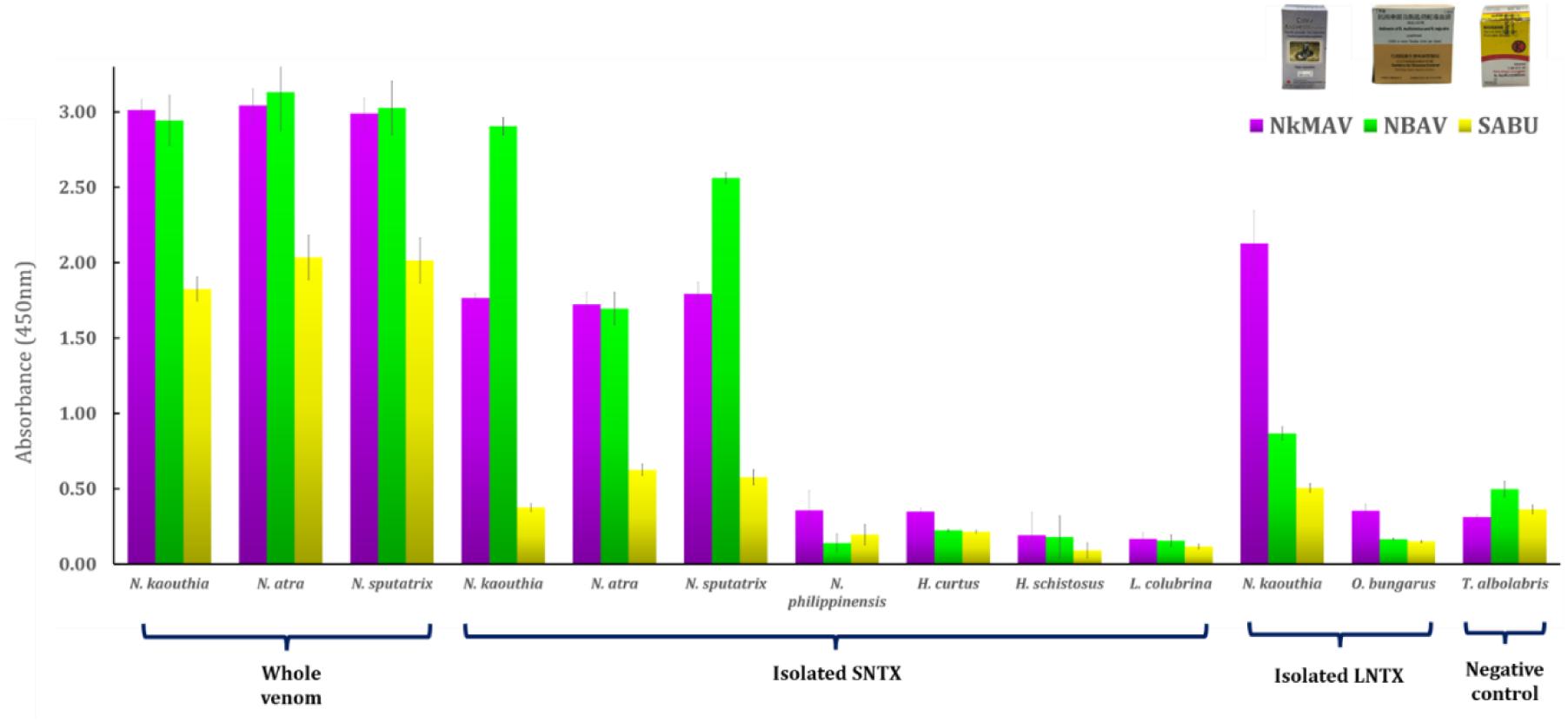
Immunoreactivity of *Naja kaouthia* Antivenom (NkMAV), Neuro Bivalent Antivenom (NBAV) and Serum Anti Bisa Ular (SABU) toward venoms and isolated alpha-neurotoxins (SNTX and LNTX) of Asian elapids. Values were expressed as means ± SEM from three independent experiments. Abbreviation of genus: *N*: *Naja*, for true cobra; *H*: *Hydrophis*, for sea snake; *L*: *Laticauda*, for sea krait; *O*: *Ophiophagus*, for King Cobra; *T*: *Trimeresurus*, for Green Pit Viper. Values were expressed as means ± SEM from three independent experiments.

### 3.3 Heatmap analysis and hierarchical clustering

Clustered heatmaps (Figure 3) summarized the immunoreactivity profiles of the four regional antivenoms. In the venom panel (Fig. 3A), PCAV displayed the strongest homologous binding to *Naja philippinensis* venom, whereas NkMAV and NBAV reacted most strongly with *N. kaouthia*, *N. atra*, and *N. sputatrix* venoms. SABU showed generally weaker responses across species. In the toxin panel (Fig. 3B), similar clustering patterns were observed: the Philippine cobra SNTX formed a distinct branch separated from the mainland Asian SNTXs, while marine SNTXs and LNTXs from *N. kaouthia* as well as *O. bungarus* grouped independently, reflecting negligible cross-recognition by all antivenoms. Overall, the clustering corroborated the geographical and species specificity of immunoreactivity patterns observed in the ELISA assays.

**Figure 3.**
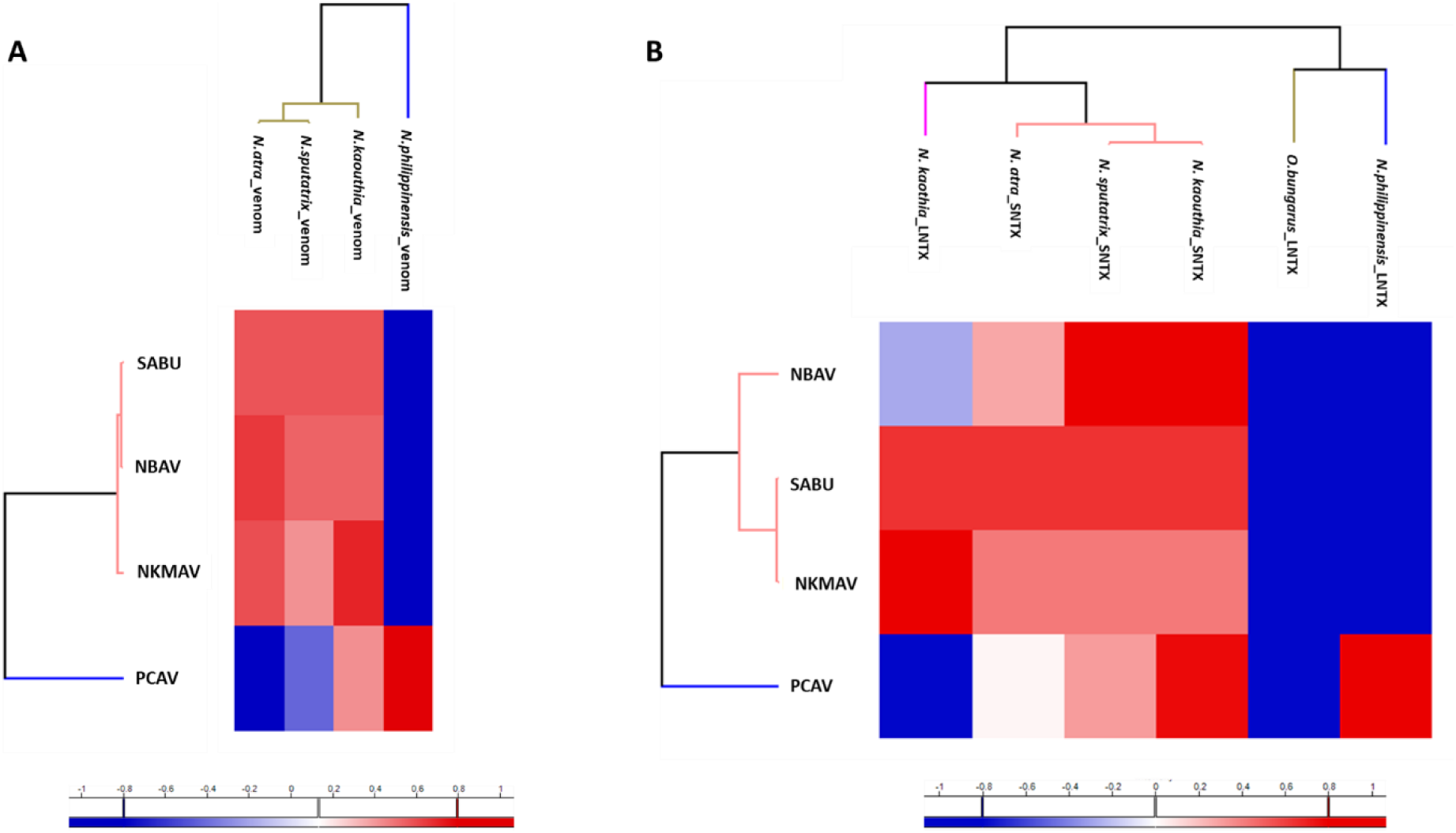
Hierarchical clustered heatmaps showing the immunoreactivity of regional antivenoms toward (A) whole cobra venoms and (B) isolated α-neurotoxins (SNTX and LNTX). Color intensity corresponds to row z-scored ELISA absorbance (A₄₅₀) values, with red indicating higher-than-average and blue lower-than-average binding for each antivenom. Abbreviations: PCAV, Philippine Cobra Antivenom; NKMAV, *Naja kaouthia* Monovalent Antivenom; NBAV, Neuro Bivalent Antivenom; SABU, Indonesian polyvalent antivenom; SNTX, short-chain α-neurotoxin; LNTX, long-chain α-neurotoxin.

### 3.4 Multiple sequence analysis of alpha-neurotoxins

Amino acid sequences of short alpha-neurotoxins tested for antivenom immunoreactivity were aligned and compared in Figure 4. All SNTX sequences reveal the conserved eight cysteine residues which result in four disulfide bridges responsible for the folding and stability of a three-finger protein structure. Residues critical for nAChR binding were indicated at the tip of loop II, while residues with high variability and non-consensus within or adjacent to the region were highlighted (in purple boxes). With numbering based on cobrotoxin of *N. atra* (UniProt: P60770, 62 amino acid residues), the 28^th^, 30^th^, 35^th^, 36^th^ and 37^th^ residues show variability and substitutions. Within the cobra species, two subsets based on the amino acid variation within this region can be recognized, as follows: W28 paired with S30 and ^35^TII^37^, or R28 paired with R30 and ^35^YRT^37^. The corresponding residues in the SNTX of two true sea snakes (*H. schistosus* and *H. curtus*) were conserved (T28, S30, ^35^TRI^37^) whereas in that of the sea krait (*L. laticauda*), the SNTX has Q28, R30, and ^35^SIT^37^ within this peptide region. The K27 residue and residues between the two motifs of the respective subsets, namely ^31^DHRG^34^, remained highly conserved within this critical region of nicotinic receptor binding.

**Figure 4.**
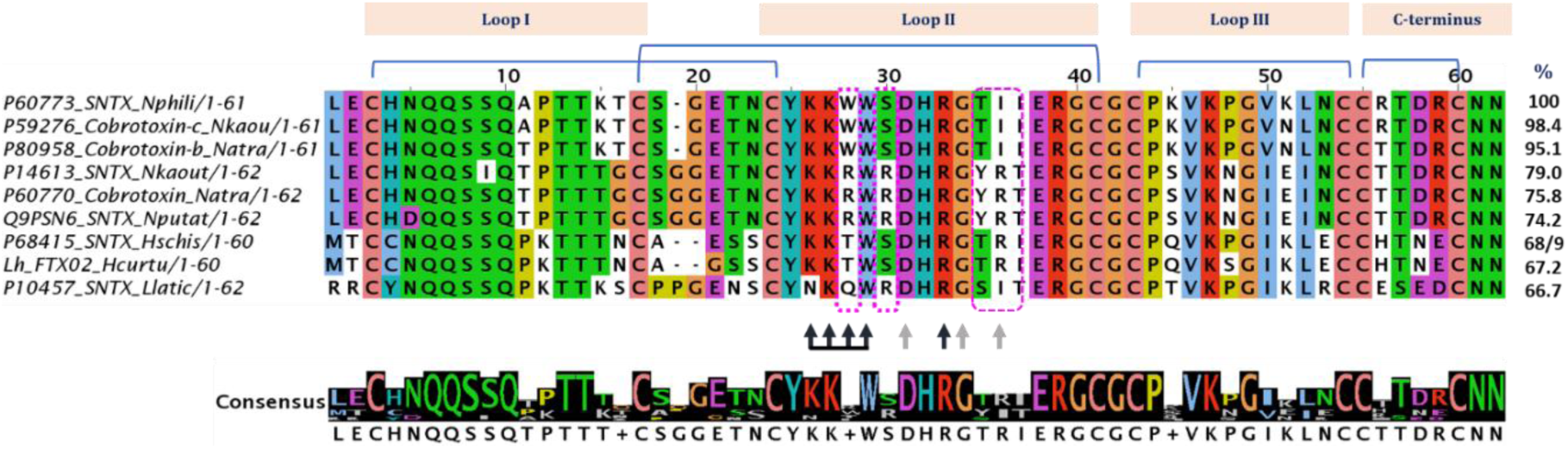
Amino acid sequences of short neurotoxins (SNTX) from elapids as used in the immunoreactivity study. Brackets indicate positions of conserved cysteine residues and disulfide bridges critical for the folding of three-finger toxins. Black and grey arrows indicate the primary and supporting residues, respectively, which are critical in binding of nicotinic receptor (nAChR). Purple boxes indicate regions with variable residues within the receptor-binding motif.

The amino acid sequences of SNTX from multiple lineages of Elapidae, including cobras (*Naja* spp., along with their subgenera), mambas (*Dendroaspis* spp.), coral snakes (*Micrurus* spp.), sea snakes (*Hydrophis* spp.), sea kraits (*Laticauda* spp.), and the Australian elapid *Suta nigriceps,* as well as the synthetic construct ScNTX were retrieved from UniProtKB and PDB databases, respectively. The sequences were further aligned for consensus analysis shown in Figure 5.

**Figure 5.**
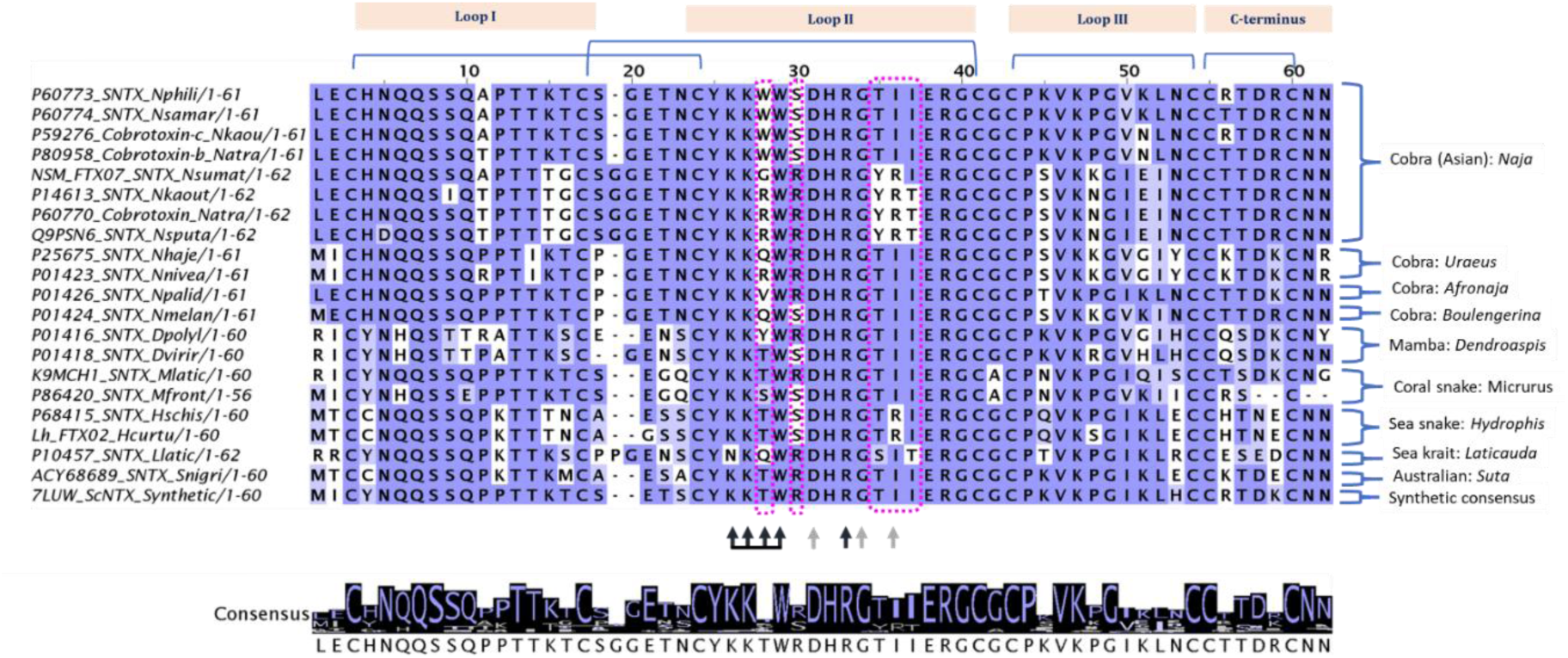
Comparison of amino acid sequences of short neurotoxins (SNTX) from diverse lineages of elapids including cobras, mamba, coral snake, sea snake, sea krait, Mallee black-back snake (Australian species, *Suta nigriceps*) and synthetic SNTX (ScNTX) which has the highest similarity to the SNTX of *S. nigriceps*. Brackets indicate positions of conserved cysteine residues and disulfide bridges critical for the folding of three-finger toxins. Black and grey arrows indicate the primary and supporting residues, respectively, critical in binding of the nicotinic receptor (nAChR). Purple boxes indicate regions with variable residues within the receptor-binding motif.

All sequences in Figure 5 retained the characteristic eight-cysteine framework, forming four disulfide bridges that stabilize the three-finger toxin structure. The sequence alignment revealed more variations (specifically, having a lower consensus) in the receptor-binding motif, particularly within loop II, where substitutions were highlighted in pink boxes. Based on numbering relative to *Naja atra* cobrotoxin (P60770, 62 amino acids), key positions of variation were primarily identified at residues 28, 30, 35, 36, and 37, consistent with Figure 4, notwithstanding greater diversity. The two subset motifs shown in Figure 4, i.e., ^28^WWS^30^----35TII^37^, and ^28^RWR^30 35^YRT^37^, appear to be consistently present in Asiatic cobras (subgenus: *Naja*) except for the variation noted in *N. sumatrana* (GWR pairing YRI). The SNTXs of African species (exhibit various combination motifs, as in Q/RWR TII in subgenus *Uraeus*, VWR----TII in subgenus *Afronaja*, and QWS TII in *Boulengerina*. The African mambas and New World coral snakes too exhibit variation in the two residues (28^th^ and 30^th^ residues based on numbering by P60770); e.g., YWR and TWS in *Dendroaspis* spp., TWR and SWS in *Micrurus* spp. while sharing the conserved TII motif. In the marine elapids, these motifs are more divergent from the terrestrial elapids, where true sea snakes (*Hydrophi*s spp.) show TWS----TRI and sea krait (*Laticauda* sp.) show QWR SIT. The synthetic ScNTX contains a TWR TII motif in this corresponding region, and exhibits the highest sequence similarity to the SNTX of *S. nigriceps*, an Australian terrestrial elapid of the Hydrophiinae subfamily in which marine elapids are nested. The overall sequence comparison also reveals conserved N-terminal sequences of SNTX specific to different clades, notable in cobras, where it is typically LECHNQQS in Asiatic cobras and, with minor substitution, MICHNQQS and MECHNQQS in African cobras. The mambas, New World coral snakes, and Hydrophiinae species show less conserved N-terminus sequence from those of the cobras while retaining the cysteine residue at the 3^rd^ residue position.

The amino acid sequences of representative short neurotoxins (SNTXs) from representative cobra species and a sea snake were aligned alongside long neurotoxins (LNTXs) from Monocled Cobra (*Naja kaouthia*), King Cobra (*Ophiophagus hannah*), and Many-banded Krait (*Bungarus multicinctus*) in Figure 6. The sequence alignment highlights conserved features essential for toxin structure and receptor interaction, as well as key regions of divergence that differentiate SNTXs from LNTXs within the three-finger protein family. All sequences maintain the three-finger loop configuration through conserved cysteine framework and disulfide-stabilized three-finger structure (brackets). Of note, the LNTXs contain an additional disulfide bond in loop II, absent in all SNTXs. The loop III and C-terminal regions of LNTXs also show greater divergence, with several charged and hydrophobic substitutions.

**Figure 6.**
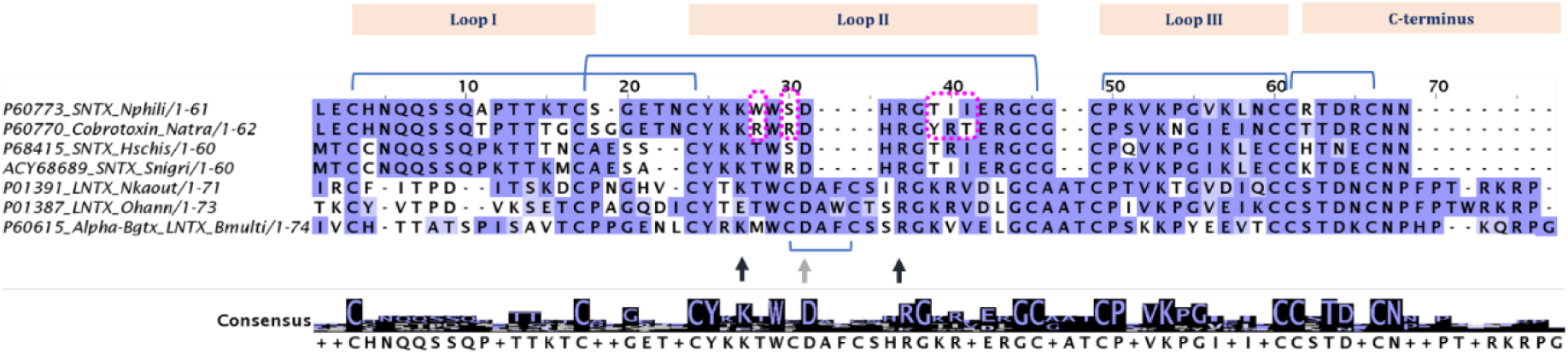
Comparison of amino acid sequences of representative short neurotoxins (SNTX) from cobras, sea snake, and long neurotoxins (LNTX) from cobra, King Cobra, and krait. Brackets indicate positions of conserved cysteine residues and disulfide bridges critical for the folding of three-finger toxins (note the additional disulfide bond in the loop II region of LNTX). Black and grey arrows indicate the primary and supporting residues, respectively, critical in binding of nicotinic receptor (nAChR). Purple boxes indicate regions with variable residues within the receptor-binding motif for the two cobra SNTX.

A low consensus region marked by differences in the receptor-binding region of Loop II is observed, especially within cobra SNTXs, which exhibit two distinct motifs, as described in Figures 3 and 4: ^28^WWS^30^ 35TII^37^ (P60773, *N. philippinesis*) and ^28^RWR^30^----^35^YRT^37^ (P60770, cobrotoxin of *N. atra*). In the hydrophiid SNTXs, the motifs are apparently more varied from cobras, e.g., substitutions resulting in TWS YRI (P68415, *H. schistosus*), and TWR----YII (ACY68680, *S. nigriceps*). LNTXs exhibit greater divergence, including additional hydrophobic and charged residues beyond position 40, as well as an additional disulfide bond, which contribute to a potentially extended receptor interaction site.

### 3.5 Phylogenetic analysis of short alpha-neurotoxins

A maximum likelihood phylogenetic tree was constructed to evaluate the evolutionary relationships of short-chain α-neurotoxins (SNTX) across representative elapid taxa, including marine, terrestrial, and arboreal lineages (Figure 6). The analysis reveals well-resolved clades that correspond to extant taxonomic groups, and illustrates the molecular diversification of SNTX sequences within and across the elapid lineages. The Hydrophiine clade, comprising the sea snakes (*H. curtus* and *H. schistosus*), branches distinctly and forms a sister group to a well-supported assemblage that includes the Australian elapid (Suta nigriceps), African mambas (Dendroaspis polylepis and D. viridis), and New World coral snakes (Micrurus frontalis and *M. laticollaris*). It is worth noting that the hydrophiid (sea snake) lineage (*Hydrophis* spp.) is positioned as an early-diverging branch among the sampled elapids, indicating a distinct evolutionary trajectory for their SNTX. This, however, does not imply that sea snakes are more “primitive“; rather, their SNTX sequence placements reflects divergence from that of a common elapid ancestor before the diversification of the terrestrial cobra-mamba-coral snake lineage. Their evolution as marine specialists suggests subsequent adaptation rather than ancestral retention of features.

The synthetic neurotoxin construct (7LUW_ScNTX), boxed in brown, was introduced into the tree and found positioned at an ancestral node closer to the branch uniting sea snakes (*Hydrophis*), Australian elapids (*Suta*), mambas (*Dendroaspis*), and coral snakes (*Micrurus*). The true cobras (*Naja*) SNTXs, on the other hand, appear as a distinct and separate clade, evolving in parallel rather than directly branching from this paraphyletic clade within which SNTXs share a higher relatedness to the synthetic construct. Also, the true cobras (genus *Naja*) were clustered into major geographic subgroups, conforming to their evolutionary divergence, which can be representative of the respective subgenera. Notably, a few SNTX sequences (marked with pink asterisks) of Asian cobras occupy basal positions within the cobra clade, and exhibit unique epitope signatures, i.e., the “WWS----TII” motif. These basal SNTX variants may represent ancestral forms from which SNTXs with the divergent epitopes “RWR YRT” evolved later within the Asiatic clade. While *N. philippinensis* and *N. samarensis* of the Philippines exhibit exclusively the ancestral form of SNTX epitopes, *N. atra* and *N. kaouthia*, along with other Asiatic cobras, evolved and prioritized the more derived forms of SNTXs. Of note, the few basal nodes within the Asian cobra clade were marked with pink asterisks that denote SNTX sequences with a unique epitope profile, essentially, carrying the “WWS----TII” motif. These likely represent ancestral molecular forms from which the more antigenically divergent SNTXs (“RWR----YRT/I” motif) of modern Southeast and East Asian cobras evolved.

The synthetic short neurotoxin construct (7LUW_ScNTX) does not group with any single natural cobra species but instead resolves near the node uniting the *Hydrophiine–Suta–Dendroaspis–Micrurus* assemblage, indicating its sequence architecture more closely resembles an inferred ancestral or consensus form of SNTX rather than a derived form typical of extant *Naja* species. This placement suggests that the synthetic construct may be functionally versatile but is “phylogenetically distinct” from the specialized SNTXs found in modern cobras.

Altogether, the tree topology supports a pattern of lineage-specific diversification of SNTX, particularly within the *Naja* genus, with antigenic divergence emerging in parallel with geographic and subgeneric separation. The integration of the synthetic sequence provides a comparative anchor to assess the evolutionary polarity of natural sequence changes, especially in regions relevant to epitope recognition and receptor binding.

### 3.6 Epitope prediction of short alpha-neurotoxins

Figure 8 shows the results of epitope predictions for four short alpha-neurotoxins (SNTXs), applying IEDB tools. The SNTXs are three wild-type cobrotoxins from *Naja kaouthia* (P32276) and *Naja atra* (P00958, P60770), and one synthetic construct (ScNTX) reported to contain the consensus sequence of epitopes from multiple SNTXs. BepiPred-2.0 predictions (Figure 7, Panel I) show the SNTX of Naja kaouthia (P32276) contained a localized epitope at Loop II, whereas those of Naja atra (P00958, P60770) had broader antigenic profiles extending beyond Loop II. The synthetic construct demonstrated epitope continuity between Loops I and II, suggesting an overlap of antigenic determinants.

**Figure 7.**
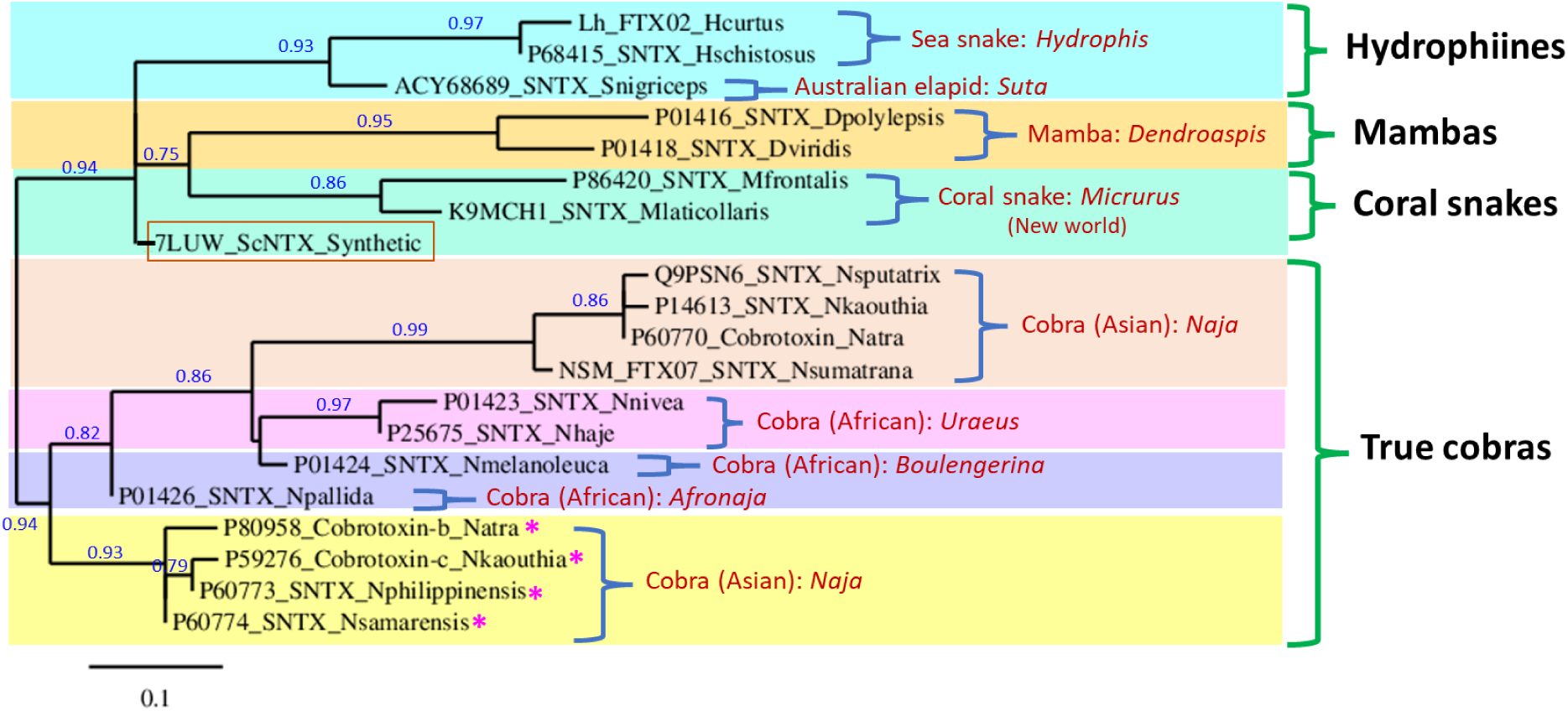
Phylogenetic tree of short neurotoxins (SNTX) from various lineages of Elapidae snakes: Hydrophiines*: Hydrophis curtus*, *Hydrophis schistosus* (both are sea snakes), *Suta nigriceps*; Mambas: *Dendroaspis polylepsis*, *Dendroaspis viridis*; Coral snakes: *Micrurus frontalis*, *Micrurus laticollaris*; True cobras: *Naja sputatrix*, *Naja kaouthia*, *Naja sumatrana*, *Naja atra*, *Naja philippinensis*, *Naja samarensis* (subgenus *Naja*); *Naja nivea*, *Naja haje* (subgenus *Uraeus*); *Naja melanoleuca* (subgenus *Boulengerina*); *Naja pallida* (subgenus *Afronaja*). Brown box contains 7LUW_ScNTX representing the synthetic SNTX based on consensus analysis reported previously [32]. Asterisks (* in pink) denote SNTX sequences at a basal position with a unique epitope which gave rise to the divergent epitopes of SNTX among the Southeast/East Asian cobras. The phylogenetic tree was constructed using the maximum likelihood method implemented in the PhyML program (v3.1/3.0 aLRT), and drawn with TreeDyn (v198.3).

**Figure 8.**
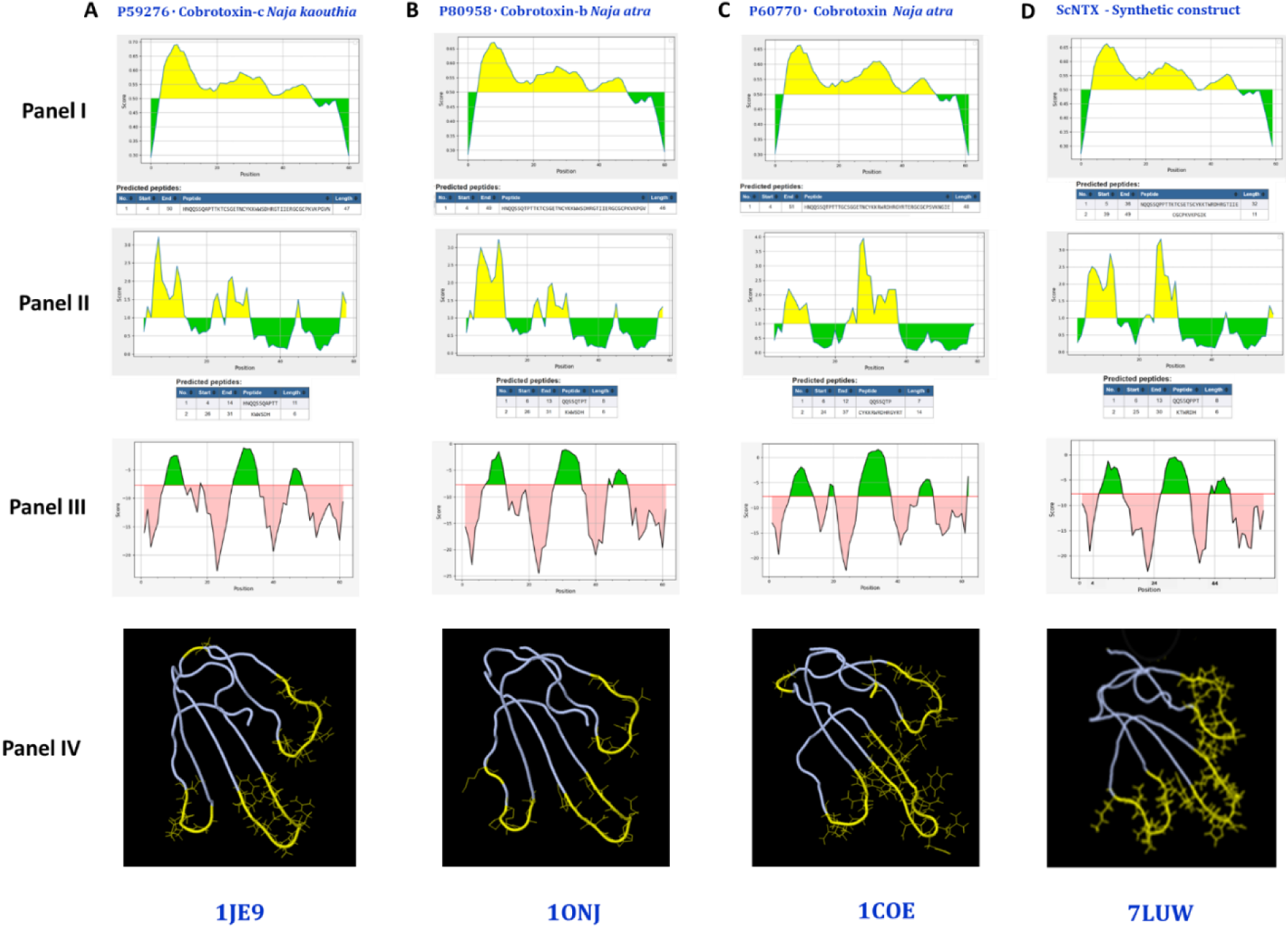
Epitope prediction for four short alpha-neurotoxins (SNTX) representing three wild types (cobrotoxins from Asiatic cobras, A: *Naja kaouthia*, B and C: *Naja atra*) and one synthetic construct (D). These SNTXs were chosen based on availability of full amino acid sequences in the UniProtKB database, and corresponding structural models available in the PDB database. Epitope predictions were conducted based on propensity scale, applying the online IEDB tools which identify B cell epitopes and peptide antigenicity using different predefined physicochemical parameters. Outputs of the analyses are illustrated according to the tools as follows: Panel I, BepiPred-2.0, score threshold = 0.5 (epitope region is colored in yellow); Panel II, Emini surface accessibility scale, score threshold = 1.0 (epitope region is colored in yellow); Panel III, DiscoTope 1.1, score threshold = −7.7 (red line); Panel IV, DiscoTope 1.1, 3d view in Jmol mode displaying structures with positive predictions (epitope residues and side chains) in yellow.

Surface-exposed antigenic sites were detected in Loops I, II, and III, with variations among the toxins (Figure 8, Panel II). The Loop III in *Naja atra* SNTXs (P00958, P60770) showed higher surface accessibility compared to that of *Naja kaouthia* SNTX (P32276). The synthetic construct maintained consistent surface exposure patterns across all loops, resembling *Naja atra* cobrotoxins. The DiscoTope 1.1 conformational epitope predictions (Figure 8, Panel III) shows regions with scores exceeding the threshold (−7.7) for epitope were mapped primarily to Loop II and Loop III, with *Naja atra* SNTXs having a more extended conformational epitope in Loop III compared to *Naja kaouthia*. The synthetic construct displayed an epitope distribution pattern similar to *Naja atra*, with higher scores in Loop II.

In Panel IV of Figure 8, structural models of the SNTXs obtained from the PDB database were analyzed for epitope localization using Jmol visualization. Predicted antigenic residues were mapped onto each structure while highlighting differences in exposure and distribution. The Loop I of *Naja kaouthia* SNTX exhibited partial burial of the predicted epitope, whereas the SNTXs of *N. atra* and the synthetic construct displayed greater solvent accessibility in this region. *N. kaouthia* SNTX contained a localized, high-scoring antigenic site at the apex of its Loop II, corresponding to the primary nicotinic acetylcholine receptor (nAChR) binding sites. *Naja atra* cobrotoxins exhibited broader antigenic regions extending along the length of Loop II while the synthetic construct integrated antigenic features from both species, with a continuous antigenic surface spanning Loop II.

## Discussion

Alpha-neurotoxins (α-NTXs), comprising long-chain (LNTX) and short-chain (SNTX) neurotoxins, are functionally conserved snake venom proteins that show potent antagonistic activity toward muscle-type nicotinic acetylcholine receptors (nAChRs) [33]. While this function appears to have evolved convergently across multiple elapid lineages as an adaptation for predation, α-NTXs are evolutionarily labile. Positive selection, driven primarily by past and perhaps ongoing evolutionary arms race, subjects these proteins to accelerated evolution [8]. This evolutionary dynamic has led to substantial sequence diversification, particularly at surface-exposed residues across elapid species, while preserving the structural framework essential for receptor binding. In comparison to SNTXs, LNTXs have traditionally received more attention owing to their higher binding affinity toward human muscle nicotinic receptor (nAChR) and thus, a “perceived” higher risk of envenomation in human [17]. Nonetheless, SNTXs are increasingly recognized for their limited immunogenic capacity (in comparison to LNTXs) [3, 19, 32], and the high abundance in the venoms of some Asian *Naja* species, particularly those from Southeast and East Asia, simply cannot be overlooked [12–14, 19]. Previous studies attributed the poor immunogenicity of SNTX to its small size (∼60–62 amino acids), an inherent factor limiting epitope display for effective immune response [20, 34]. Beyond the molecular size of the SNTX, in this study, we showed that SNTXs display subtle sequence-level variations that may critically influence their antigenicity and thus, impacting the effectiveness of antivenom therapy. As revealed in a series of Asiatic cobra venomics, there is a transition from LNTX to SNTX-dominant phenotype as cobras diversified from the west to the east, with the easternmost dispersal lineage of *Naja* spp., represented by *N. philippinensis* and *N. samarensis* of the Philippines, characterized by extremely lethal venoms containing exceptionally high abundance of SNTXs (44-66%) but no LNTXs [12, 13]. The same is true for *N. atra*, albeit with a lower abundance of SNTX (∼10%) and lower lethality [14, 23]. Intriguingly, the Philippine spitting cobra venom could not be effectively cross-neutralized by antivenom raised against *N. atra*, and vice versa, suggesting antigenic variation amongst the two (unpublished data). On the other hand, the venoms of other Asian *Naja* spp. had lower abundances of SNTXs, typically ranging from 1–10% of total venom protein (Table 1). The remarkable variation in both abundance and epitope composition of SNTXs between the Philippine species and other Asian cobras may explain the reduced immunoreactivity of PCAV toward venoms and isolated SNTX of these Southeast Asian cobras.

**Table 1:**
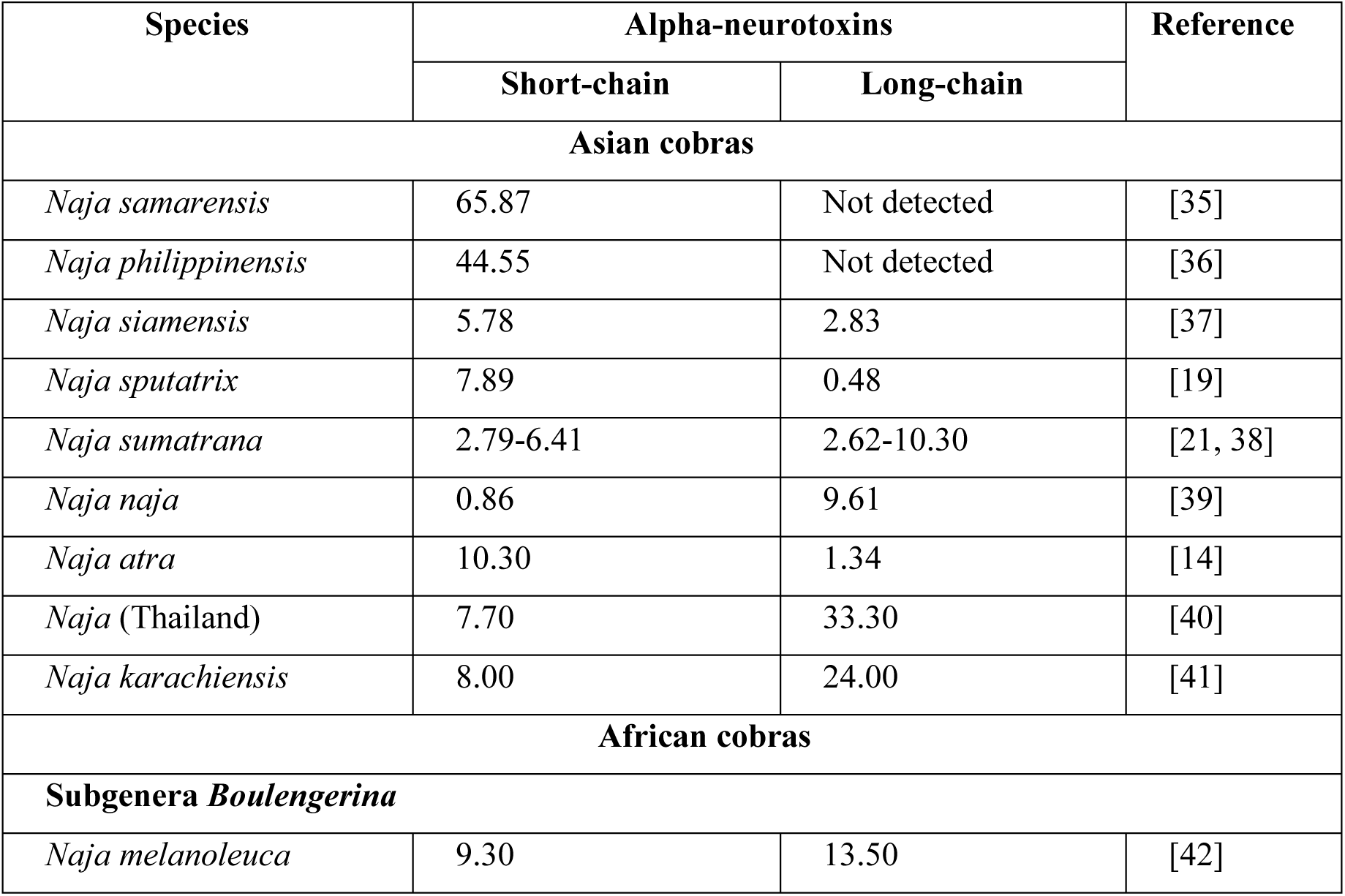
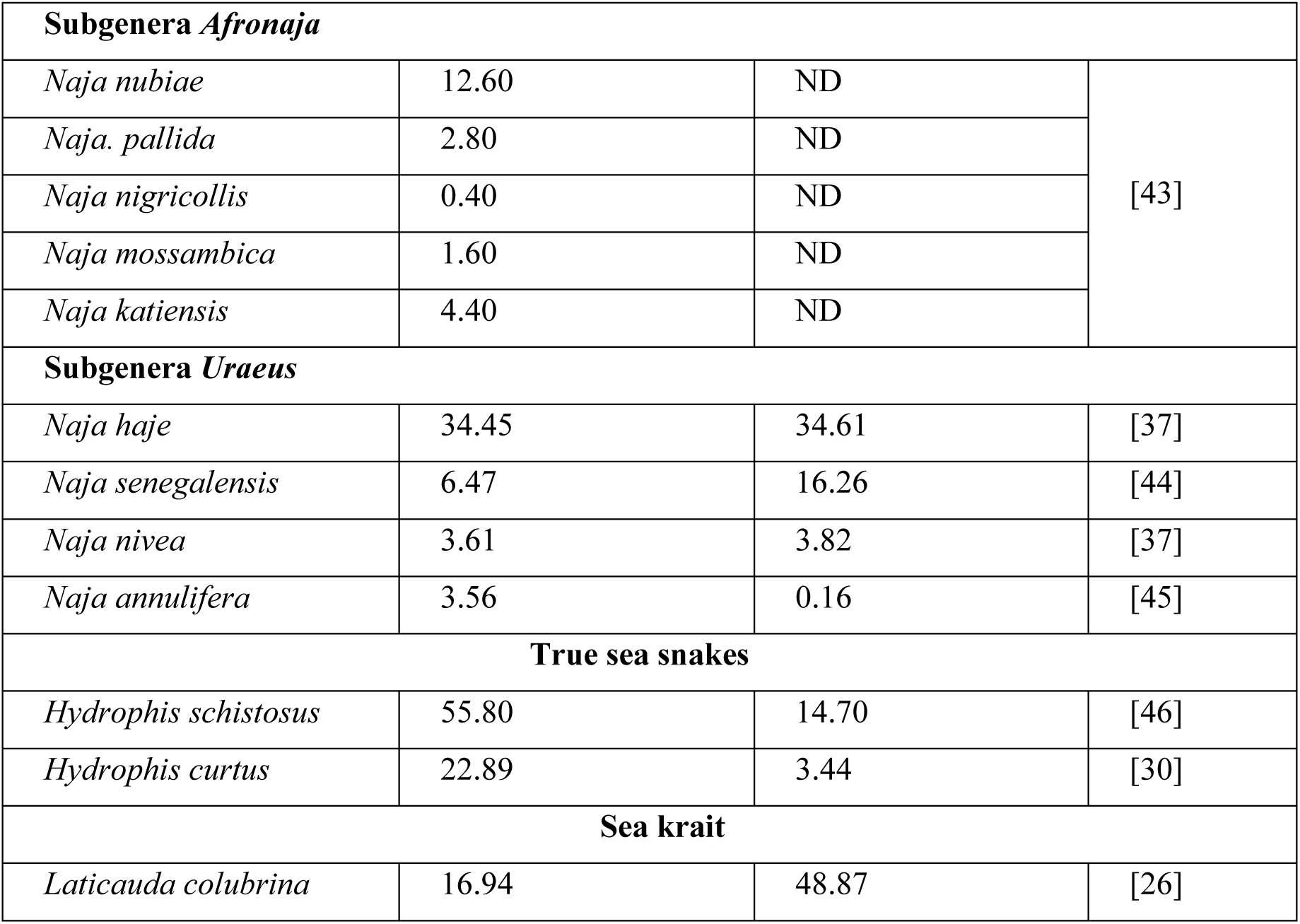
Composition of short- and long-chain alpha-neurotoxins from representative Afro-Asian cobras (*Naja* spp.), and marine elapid species.

| Species | Alpha-neurotoxins |  | Reference |
| --- | --- | --- | --- |
|  | Short-chain | Long-chain |  |
| Asian cobras |  |  |  |
| *Naja samarensis* | 65.87 | Not detected | [35] |
| *Naja philippinensis* | 44.55 | Not detected | [36] |
| *Naja siamensis* | 5.78 | 2.83 | [37] |
| *Naja sputatrix* | 7.89 | 0.48 | [19] |
| *Naja sumatrana* | 2.79-6.41 | 2.62-10.30 | [21, 38] |
| *Naja naja* | 0.86 | 9.61 | [39] |
| *Naja atra* | 10.30 | 1.34 | [14] |
| *Naja* (Thailand) | 7.70 | 33.30 | [40] |
| *Naja karachiensis* | 8.00 | 24.00 | [41] |
| African cobras |  |  |  |
| Subgenera *Boulengerina* |  |  |  |
| *Naja melanoleuca* | 9.30 | 13.50 | [42] |

| Subgenera *Afronaja* |  |  |  |
| --- | --- | --- | --- |
| *Naja nubiae* | 12.60 | ND | [43] |
| *Naja. pallida* | 2.80 | ND |  |
| *Naja nigricollis* | 0.40 | ND |  |
| *Naja mossambica* | 1.60 | ND |  |
| *Naja katiensis* | 4.40 | ND |  |
| Subgenera *Uraeus* |  |  |  |
| *Naja haje* | 34.45 | 34.61 | [37] |
| *Naja senegalensis* | 6.47 | 16.26 | [44] |
| *Naja nivea* | 3.61 | 3.82 | [37] |
| *Naja annulifera* | 3.56 | 0.16 | [45] |
| True sea snakes |  |  |  |
| *Hydrophis schistosus* | 55.80 | 14.70 | [46] |
| *Hydrophis curtus* | 22.89 | 3.44 | [30] |
| Sea krait |  |  |  |
| *Laticauda colubrina* | 16.94 | 48.87 | [26] |

Based on the previous study of the Philippine Cobra Antivenom (PCAV) neutralization against the venoms of *Naja samarensis* and *Naja philippinensis* [12, 13], the present immunoreactivity findings confirm that these far-eastern, insular cobras endemic to the Philippines possess SNTXs with distinct antigenic profiles, differentiating them from other Southeast Asian cobras. Notably, PCAV showed markedly reduced cross-reactivity toward SNTXs from non-Philippine cobra species, following the order: *N. kaouthia* > *N. sputatrix* > *N. atra*. This pattern may reflect historical biogeographic dispersal, in which the Philippine cobra species could have descended from an ancestral lineage in continental Asia that radiated southward and eastward, with *N. kaouthia* and *N. sputatrix* representing intermediate lineages, while *N. atra* diverged along an eastward trajectory into mainland China approximately 1–2 Mya, and subsequently reached Taiwan during the Late Pleistocene glacial periods (∼20,000–70,000 years ago) before the submersion of the land bridge. In Taiwan’s insular and montane environments, further ecological adaptation in *N. atra* might have resulted in a venom phenotype that includes SNTXs with antigenic features that are much divergent from their Philippine counterparts. In fact, even within the mountainous island of Taiwan, the central ridge has possibly been a biogeographical barrier, resulting in two variable phenotypes between the eastern and western *N. atra* populations, as reported in both their morphology and venom compositions [14, 47]. The present study also confirmed the very limited cross-reactivity of PCAV toward SNTXs of marine elapids, including sea kraits and sea snakes, as well as LNTXs from both *N. kaouthia* (a true cobra) and *O. bungarus* (King Cobra) in Southeast Asia due to the greater variation in sequences and epitopes.

We also further evaluated the immunorecognition of αNTXs by other cobra antivenoms available in the region: *Naja koouthia* Monovalent Antivenom (NkMAV, Thailand), Neuro Bivalent Antivenom (NBAV, Taiwan), and Serum Anti Bisa Ular (SABU, Indonesia). NkMAV showed significantly higher immunological binding activity among the antivenoms toward LNTXs of *N. kaouthia*, which is intuitively appropriate as the Thai *N. kaouthia* venom is used as its immunizing venom. Furthermore, in comparison to various other southeast Asian cobras including its own species from other geographical locales, the Thai *N. kaouthia* venom contains the highest abundance of LNTXs (35% of total venom protein) [15]. Hypothetically, NkMAV is a LNTX-targeting antivenom for cobra species within the region, e.g., *N. sumatrana,* which also shares a similar LNTX-dominant venom phenotype. Indeed, NkMAV is used clinically as the antivenom treatment in Malaysia where *N. kaouthia* and *N. sumatrana* are both found [48], with success to some extent. However, NkMAV is not effective and not indicated for envenoming by the King Cobra (*O. hannah*) in spite of its LNTX-dominant venom phenotype, since King Cobra’s and *Naja* cobra’s LNTXs are evolutionarily and antigenically diverged [49]. While exhibiting high immunoreactivity toward the *N. kaouthia* LNTX, NkMAV consistently showed a lower immunoreactivity by at least 30% toward the SNTXs of its own and heterologous *N. sputatrix* as well as *N. atra*, implying that an “LαNTX-targeting” antivenom, as proposed above, has weak binding activity for SNTX of these cobras. On the other hand, NBAV, raised against Taiwan Cobra venom which contains only SNTX as its sole alpha-neurotoxin [14, 23], exhibited higher binding activities toward these SNTXs. Meanwhile, NBAV’s binding activity toward the LNTX is, unsurprisingly, significantly lower than NkMAV. The finding further supports that LNTX and SNTX are indeed antigenically variable, with the latter (SNTX) having a lower immunogenicity than LNTX [20]. The Indonesian antivenom SABU, unfortunately, had the overall lowest immunoreactivity toward all alpha-neurotoxins, a finding consistent with its polyvalent (trivalent) nature (having the smallest portion of SNTX from *N. sputatrix*), a lower content of immunoglobulins, and a lower immunological binding activity as well as neutralization efficacy against various venoms [50]. Finally—a finding that prompted the question of antigenic divergence in the SNTX of southeast Asian cobras: all three antivenoms (NkMAV, NBAV, SABU) lack immunoreactivity toward the SNTXs of *N. philippinensis* and marine elapids, highlighting species-specific and biogeographically influenced epitope variation in the cobra SNTXs.

The heatmap visualization highlights the antigenic divergence between the Philippine and other Asian cobra neurotoxins. The strong homologous binding of PCAV to *N. philippinensis* SNTX contrasts with the limited cross-recognition by other regional antivenoms, supporting the notion that short-chain α-neurotoxins have evolved with geographically distinct antigenic epitopes. NkMAV and NBAV clustered closely, indicating shared recognition of major SNTXs from *N. kaouthia*, *N. atra*, and *N. sputatrix*, whereas all antivenoms showed weak recognition of long-chain α-neurotoxins, particularly *O. hannah* LNTX. These findings suggest that current immunization mixtures emphasize SNTX antigens of regional cobras, leading to restricted cross-neutralization, and reinforce the need to incorporate representative antigens from both SNTX motif lineages to broaden antivenom coverage.

Assuming the antigenicity divergence arises from the epitope variability of SNTXs, we further aligned and analyzed the SNTX sequences to understand if any variation in the sequence motifs might serve as drivers of antigenic subclasses. The sequence analysis reveals two dominant loop II motifs in Asian cobras: “^28^WWS----TII^37^” and “^28^RWR----YRT^37^”, between which the sequence “^31^DHRD^34^” remained highly conserved across various lineages. *N. philippinensis* and *N. samarensis* expressed SNTXs with only the “^28^WWS TII^37^” motif, whereas *N. atra* (Taiwan) and *N. kaouthia* (Thailand) express two forms of SNTXs, each contains “^28^WWS----TII^37^” and “^28^RWT----YRT^37^”, respectively, with the latter form “^28^RWT----YRT^37^” being the majority. The *N. sputatrix* (Indonesia) SNTX contains only the “^28^RWT----TRT^37^” form of SNTX, while that of *N. sumatrana* (Malaysia) in this study revealed a slight variation in this motif as “^28^GWR YRI^37^”. The 28^th^ glutamine (^28^G) or arginine (^28^R) residue has a different property from the tryptophan residue (^28^W) which contains a large hydrophobic, double-ring aromatic side chain; hence, the *N. sumatrana* SNTX should appear similar structurally to the “^28^RWT YRT^37^” motif of *N. sputatrix*, which is also shared between *N. kaouthia* (Thailand) and *N. atra* (Taiwan) as their major form of SNTX. These dichotomous motifs agree with the variable immunoreactivity of PCAV that binds effectively to SNTX containing the motif “^28^WWS----TII^37^” but not “^28^RWT----YRT^37^”, while the reverse is true for NBAV, NKMAV, and SABU (i.e., binding effectively to SNTX having the motif ^28^RWT----YRT^37^” but not “^28^WWS TII^37^”. The finding suggests an epitope-level divergence between the two motifs in these cobra SNTXs despite sharing relatively close phylogenetic relatedness and biogeographical proximity in Southeast Asia and East Asia. On the other hand, the SNTXs of African cobras vary from those of the Asiatic species to a much greater extent, whereby the 28^th^ amino acid residue is substituted by glutamine (Q) and valine (V) while retaining the conserved region of ^29^W and ^30^R residues within the motif. The African mambas, New World coral snakes, and hydrophiid species have an even more variable combinations of the motif (^28^XWX^30^) within this critical region of loop II, further indicating that although this key functional site remain conserved for receptor binding (Hegde et al., 2010), surface-exposed residues in the vicinity of the loop can mutate and diverge antigenically. Ostensibly, the mutation begat variable epitopes which brought advantage in a predator-prey arms-race, as these cobras of different species adapt to distinct ecological niches throughout Asia and Africa. Interestingly, the synthetic construct (ScNTX) harbors the motifs of “^28^TWR^30^” and “^35^TII^37^” (along with the all-conserved “^31^DHRG^34^” motif) in its loop II, which are, by sequence comparison, highly similar (nearing 100%) to the SNTX of an exotic and endemic terrestrial elapid *Suta nigriceps* from Australia. Its “^28^TWR^30^” motif, in comparison to the abovementioned “^28^RWR^30^” and “^28^WWS^30^” motifs of Asiatic cobras’ SNTXs, deserves a note of distinction in terms of their structure and antigenicity. Both the constructed “^28^TWR^30^” and the native “^28^RWR^30^” contain tryptophan (W) and arginine (R) at positions 29 and 30, with only one residue difference, *i.e.*, threonine (T) *vs.* arginine (R) at position 28. Threonine is small and polar, while arginine is larger and basic. Still, the difference is not as drastic as introducing a tryptophan (W) as in the case of the “^28^WWS^30^” motif, which contains two bulky tryptophan residues—this could alter the local conformation significantly. Also, its 30^th^ residue is substituted by serine (S), an uncharged polar residue, unlike the basic (typically also more antigenic) arginine (R) at position 30. Hence, both “^28^TWR^30^” and “^28^RWR^30^” should have a higher potential to form a similar antigenic surface. At the same time, the “^28^WWS^30^” motif contributes to a surface of higher hydrophobicity and lower charge, leading to a lower degree of antigenic similarity to the other two.

On the other hand, the secondary variable motif “^35^TII^37^” is conserved with that of the SNTXs carrying “^28^WWS^30^” motif (notably, in *N. philippinensis* and *N. samarensis*), and those of African cobra species. Comparing between “^35^TII^37^” and “^35^YRT^37^” motifs (the latter is paired with the “^28^RWR^30^” motif, predominantly found in *N. kaouthia*, *N. sputatrix* and *N. atra*), the ^35^TII^37^ motif likely contributes to protein core structure or membrane interactions rather than immunorecognition, as this relatively compact region is mostly hydrophobic with low surface exposure for antibody accessibility. Further sequence comparison between representative short neurotoxins (SNTXs) from *Naja philippinensis*, *N. atra*, *H. curtus*, and *S. nigriceps*, and long neurotoxins (LNTXs) from *N. kaouthia*, *O. hannah* (*O. bungarus*), and *B. multicinctus* revealed greater divergence in residues flanking the critical sites. Notably, in the LNTXs analyzed, the 30^th^ residue is replaced by a cysteine, creating motifs such as ^28^TWC^30^ or ^28^MWC^30^. This substitution enables the formation of an additional loop, stabilized by a disulfide bond between the cysteine residues at positions 30 and 35. This innovative structural role has a functional implication in which a conformational constraint is introduced to stabilize the local structure of loop II, modulating the surface topology that possibly enhances binding affinity to muscle-type nicotinic acetylcholine receptors (nAChRs) through greater hydrophobic and electrostatic contacts [51].

Overall, the notable variation at positions 28, 30, and 35 occurring within structurally tolerant regions of the cobra alpha-neutrotoxins is in line with the “Three-Fingered RAVER” theory (rapid antigenic variation in exposed residues) in the three-finger toxin superfamily [8]. The variation reflects accelerated evolution as an adaptive strategy to diversify antigenicity without compromising receptor binding or venom potency. Considering the high degree of evolvability in this class of toxins, the pursuit of a universal anti-SNTX antivenom that is ubiquitously effective against all SNTX types deserves further and thorough investigation, including the search for non-antibody-based alternatives such as small-molecule inhibitors and *de novo* designed mini-binders of venom proteins [52, 53]. Nonetheless, antivenoms will likely remain the cornerstone of therapy for the foreseeable future. Therefore, optimizing antivenom formulations should involve incorporating a balanced repertoire of anti-LNTX and anti-SNTX antibodies, taking into account not only the structural subtypes of 3FTx toxins but also their relative abundance within the venom proteome, as well as potential epitope variations that may occur even among structurally conserved toxins across geographically distinct species.

## Conclusion

This study offers a new perspective on the divergence of cobra venom short-chain α-neurotoxins (SNTXs), which exhibit significant antigenic variability despite structural conservation. The Philippine spitting cobras (*N. philippinensis* and *N. samarensis*) express SNTXs with an ancestral loop II motif (^28^WWS–TII^37^), which are strongly recognized by PCAV but poorly by other regional antivenoms. In contrast, Asiatic cobras such as *N. kaouthia*, *N. atra*, and *N. sputatrix* predominantly express a divergent and more derived form of corresponding motif (^28^RWR–YRT^37^), which may explain the limited cross-neutralization by PCAV. Conversely, antivenoms raised against the ^28^RWR–YRT^37^ type of SNTX (predominant forms in *N. kaouthia*, *N. atra*, *N. sputatrix*) neither bound nor neutralized the ^28^WWS–TII^37^ type of SNTX from the Philippine cobras. These findings underscore that antigenicity differences at key epitope residues, rather than size constraints alone, contribute to the poor antigenicity of SNTXs. From a clinical perspective, this divergence highlights the challenges of developing a universal anti-cobra antivenom. Together, our findings highlight that SNTX antigenicity is shaped by discrete sequence motifs, which define regional immunoreactivity barriers. Rational antivenom development in the future should integrate these antigenic subtypes alongside LNTX to broaden the cross-species efficacy of antivenom.

## Acknowledgment

The authors would like to acknowledge funding support from the National Science and Technology Council (NSTC) of Taiwan (114-2311-B-007-005), funding for international collaboration from the National Tsing Hua University (114QI010E1), and the special research grant of Universiti Malaya, Malaysia (BKS003-2020).

